# Identification of Cellular Signatures Associated with Chinese Hamster Ovary (CHO) Cell Adaptation for Secretion of Antibodies

**DOI:** 10.1101/2024.09.30.614046

**Authors:** Ying Bai, Ivan Domenech Mercadé, Ramy Elgendy, Giulia Lambiase, Sew Yeu Peak-Chew, Catarina Franco, Steven W Wingett, Tim J Stevens, Luigi Grassi, Noah Hitchcock, Cristina Sayago Ferreira, Diane Hatton, Elizabeth A. Miller, Rajesh K. Mistry

## Abstract

The secretory capacity of Chinese hamster ovary (CHO) cells remains a fundamental bottleneck in the manufacturing of protein-based therapeutics. Unconventional biological drugs with complex structures and processing requirements are particularly problematic. Although engineered vector DNA elements can achieve rapid and high-level therapeutic protein production, a high metabolic and protein folding burden is imposed on the host cell. Cellular adaptations to these conditions include differential gene expression profiles that can in turn influence the productivity and quality control of recombinant proteins. In this study, we used quantitative transcriptomics and proteomics analyses to investigate how biological pathways change with antibody titre. Gene and protein expression profiles of CHO pools and clones producing a panel of different monoclonal and bispecific antibodies were analysed during fed-batch production. Antibody-expressing CHO pools were heterogeneous, resulting in few discernible genetic signatures. Clonal lines derived from these pools, selected for high and low production, yielded a small number of differentially expressed proteins that correlated with productivity and were shared across biotherapeutics. However, the dominant feature associated with higher protein production was transgene copy number and resulting mRNA expression level. Moreover, variability between clones suggested that the process of cellular adaptation is variable with diverse cellular changes associated with individual adaptation events.

## Introduction

The development and manufacture of biotherapeutics exploits the biosynthetic secretory capacity of professional secretory cell lines such as Chinese hamster ovary (CHO) cells (Hansen et al., 2017; Zhou et al., 2018). The ability of CHO cells to accommodate the expression of a diverse portfolio of protein-based therapeutics, including hormones (Sinegubova et al., 2024), clotting factors (Kovnir et al., 2018), monoclonal antibodies (Kim et al., 2012; Majumdar et al., 2024) and multi-specific formats such as bispecific antibodies (Wang et al., 2019), owes in part to their genetic tractability and adaptability.

It has long been established that heterologous genetic elements used in the expression vector are a major determinant of biotherapeutic protein expression in CHO cells, and thus have a strong bearing on overall productivity (Dong et al., 2019; Ho et al., 2015; Kim et al., 2011; Li et al., 2018; Neville et al., 2017; Nguyen et al., 2019). Commercial vectors are often engineered to drive constitutively high levels of recombinant protein transcription that consequently leads to a high protein folding burden and a strong selection pressure toward host cell adaptation (Davies et al., 2011; Ong et al., 2019; Zou et al., 2018). These sometimes-extreme adaptations arise from genetic changes involving the differential expression of genes that are critical for processes such as endoplasmic reticulum associated protein folding (Chung et al., 2004; Fatima et al., 2021; O’Rourke et al., 2019), quality control and degradation (Mathias et al., 2020) and vesicular trafficking (Peng et al., 2011; Peng & Fussenegger, 2009; Peng et al., 2010).

Despite success in developing genetic elements that promote protein production, and the adaptive capacity of CHO cells, productivity outcomes and associated adaptations remain somewhat variable. Some “easy to express” (ETE) biopharmaceuticals engender extremely high levels of productivity that necessitate metabolic changes to meet the desired cost of production, placing a further burden on the secretory pathway (Yao et al., 2021). As such, several studies have identified metabolic signatures associated with improved antibody titre including yeast cytosolic pyruvate carboxylase (PYC2) (Fogolin et al., 2004; Gupta et al., 2017), mitochondrial malate dehydrogenase II (MDH II) (Chong et al., 2010) and the fructose transporter gene, Slc2a5 (Inoue et al., 2010; Wilkens & Gerdtzen, 2011). By contrast, many novel format antibodies are engineered to contain non-native domains that can lead to protein misfolding and aggregation that require further cellular adaptations to help resolve (Brinkmann & Kontermann, 2017; Chiu & Gilliland, 2016; Feige et al., 2010). Indeed, previous work focusing on such “difficult-to-express" (DTE) proteins reveals genetic adaptations involved in apoptosis (Li et al., 2022) and protein degradation processes such as endoplasmic-reticulum-associated protein degradation (ERAD) (Mathias et al., 2020) as important for DTE antibody expression. Moreover, CHO cell engineering strategies focused on directing host cell adaptation towards desired phenotypes such as cold adaptation (Syddall et al., 2024), improved cellular redox (Mistry et al., 2021) and selection for enhanced mitochondrial performance (Chakrabarti et al., 2022) have proved effective for boosting productivity of ETE and DTE antibodies (Chong et al., 2010; Fogolin et al., 2004; Gupta et al., 2017; Inoue et al., 2010; Wilkens & Gerdtzen, 2011; Yao et al., 2021).

Given the bespoke and often complex folding and trafficking needs for biotherapeutics, it is not surprising that successful production can be unpredictable. It would be commercially advantageous to overcome such stochasticity and identify cellular signatures that consistently associate with high productivity. Such gene expression changes have previously been characterized on a case-by-case basis. Here, we sought to survey a larger panel of cell lines expressing a diverse set of antibody-based biotherapeutics to understand host cell genetic adaptations that occur in the setting of high titre secretion and in response to different antibody formats. To do this we compared heterogeneous pooled lines and clonal cell lines expressing ETE antibodies with multiple lines expressing more complex DTE modalities, which in some instances displayed aberrant product quality attributes. This analysis revealed many adaptive changes in the transcriptome and proteome of cells selected for high and low productivity. A surprisingly small number of individual hits were identified as shared across productivity outcomes and might therefore be key drivers of enhanced antibody production. However, there was a distinct lack of conserved, correlated biological “signatures” associated with secretion phenotypes. Our findings suggest that cellular adaptations are difficult to predict and therefore remain a challenge for CHO cell-mediated biotherapeutic production.

## Methods

### Cell lines and Culture Conditions

AstraZeneca proprietary CHO host cells were maintained in CD-CHO medium (Life Technologies) supplemented with 6CmM L-glutamine (Life technologies) and sub-cultured every 3 - 4 days at a seeding density of 0.2 - 0.3 ×C10^6^Ccells/ml. Stably transfected CHO cell pools and clones were routinely maintained in AstraZeneca proprietary medium. Routine culture for all cell lines was performed at 36.5°C in vented, non-baffled Erlenmeyer flasks (Corning, Amsterdam, The Netherlands), in 5 % CO_2_ maintained at 140 rpm. Cells were sub-cultured every 3 - 4 days at a seeding density of 0.3 - 0.4C×C10^6^Ccells/ml. Viable cell density and cell viability were assessed using a Vi-Cell XR automated cell counter and viability analyser (Beckman Coulter, High Wycombe, UK).

### Vector Construction

Immunoglobulin heavy chain (HC) and light chain (LC - kappa) encoding DNA sequences were synthesised (Thermo Fisher, UK) and cloned into a dual chain expression vector with the human cytomegalovirus (CMV) promoter driving expression of HC and LC. The expression vector also contained a glutamine synthetase (GS) selectable marker gene. A schematic map of a linear plasmid vector used in this study is illustrated in Figure 1B. All vectors were constructed based on standard cloning methods. The cloned sequences were confirmed by DNA sequencing. The MonoAb1- and MonoAb2-expressing plasmids were modified from BisAb1 and BisAb2 expression vectors respectively by removing the single-chain variable fragment (scFv) sequence by standard restriction enzyme digestion and ligation. A no IgG control vector was also generated which did not contain HC or LCs. Transfection-grade plasmid DNA was purified using the QIAGEN maxi prep plus kit (Qiagen, Manchester, UK) and subsequently linearised via restriction enzyme digest.

**Figure 1:**
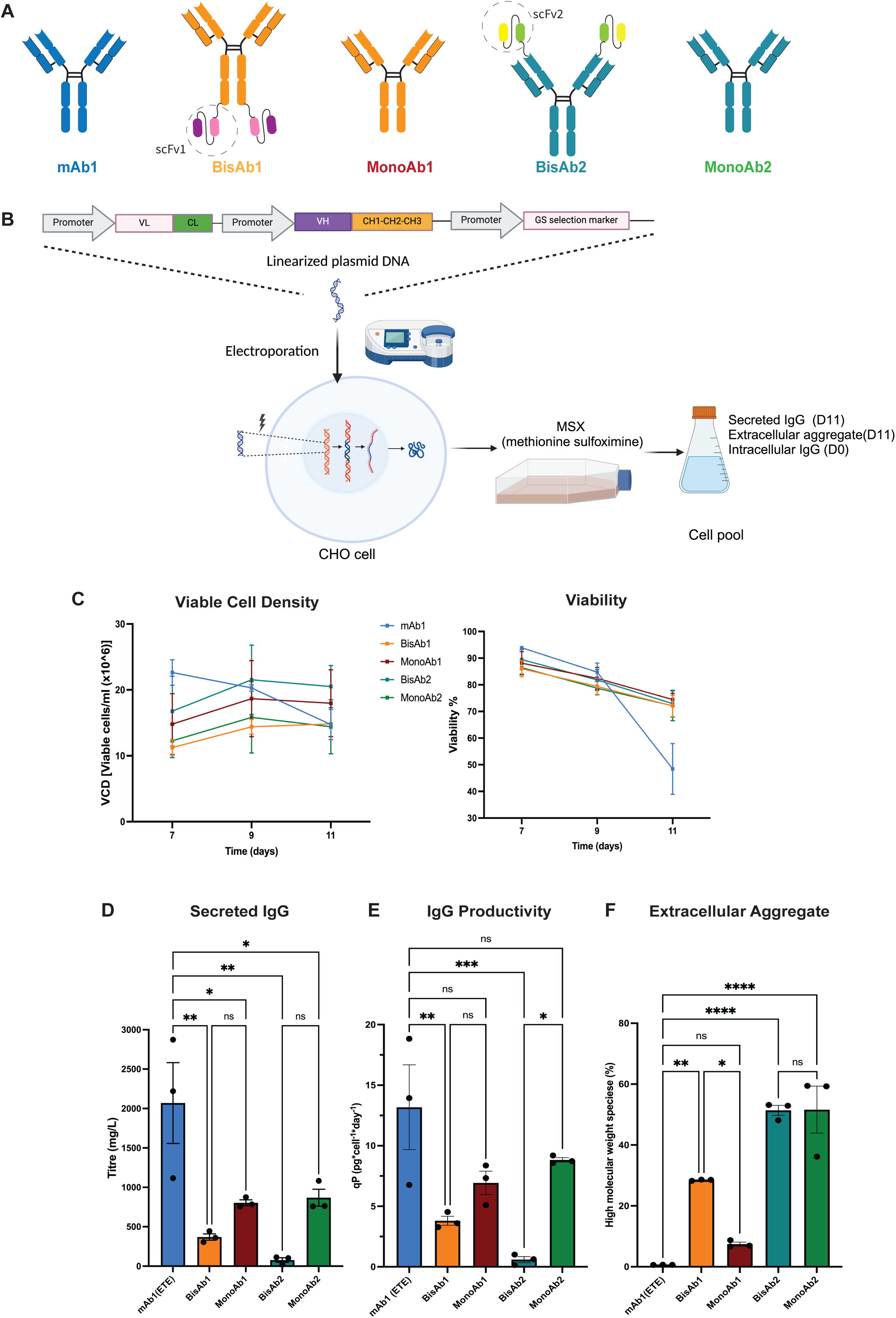
Generation and Characterisation of Stable Cell Pools Expressing Different Antibody Format Biotherapeutics. **A:** Diagrams detailing the structure of mAb1 (Monoclonal antibody 1) in blue, BisAb1 (Bispecific antibody 1), MonoAb1 (Monovalent control for BisAb1) in orange, BisAb2 (Bispecific antibody 2) and MonoAb2 (Monovalent control for BisAb2) in teal. BisAb1 and BisAb2 contain single chain variable fragment (ScFv1/2) domains that permit bivalent antigen recognition which were removed to create monovalent controls for each bispecific antibody. **B:** A schematic detailing stable cell pool generation. **C:** Characterisation of stable pool parameters over an 11-day fed-batch process. Viable cell number and viability measured at D7, D9 and D11 are presented for each cell pool. **D:** Secreted IgG titre measured at D11. **E**: Cell specific productivity for each pool. **F:** Biotherapeutic high molecular weight species percentage for each pool. Data presented as mean ± standard deviation (SD) from N=3 biological replicates, P value *<0.05. **<0.01, *****<0.0001. Statistical analysis was performed using One-way ANOVA with multiple comparisons corrected by Tukey post hoc analysis.

### Stable Pool Generation

Stable transfectant pools expressing no IgG control, mAb1, MonoAb1, MonoAb2, BisAb1 and BisAb2 were generated using the Amaxa^TM^ Nucleofector^TM^ system (Lonza, Basel, Switzerland) with the Cell Line Nucleofector® Kit V (Lonza) as per manufacturer’s instructions. In brief, the linearised DNA was resuspended in cell-line Nucleofection solution V, this mixture was then combined with AstraZeneca-proprietary CHO host cells; the cell-DNA mix was nucleofected before the transfected cells were combined with pre-warmed CD CHO medium in a T175cm^2^ flask and incubated at 36.5°C static incubator (5% CO_2_, 80% humidity). Next day, methionine sulfoximine (MSX) solution was added to T175 cm^2^ flask to a final concentration of 50 µM. Cell viability and viable cell density (VCD) were monitored on Vi-Cell XR automated cell counter (Beckman Coulter, High Wycombe, UK) until cells recovered.

### Clonal Cell Generation

Prior to single-cell cloning, 384-well plates (Corning, Cat. No. 3680) were filled with CD-CHO medium (Thermo Fisher Scientific,UK) supplemented with 50 μM MSX (Sigma-Aldrich) and 1x diluted Solentim InstiGRO^TM^ CHO (RS-1105, Advanced Instruments) using a Multidrop Combi reagent dispenser (ThermoFisher Scientific, UK). For single-cell cloning, each CHO pool to be cloned was sorted by depositing single cells into individual wells of 4-6 pre-filled 384-well plates using the f.sight single-cell printer (cytena, Germany). Once printing was completed, plates were centrifuged for 5 min at 2000 x g. All plates were then incubated at 36.5°C in a humidified incubator with 6 % CO_2_ for 11 - 14 days before cell confluency was measured on a Cellavista imager. Clones with confluence over 20% were expanded into flat bottom 96-well plates (Corning,Cat. No. 3598) and then into a 96-well 2-ml Masterblock plate (Greiner bio-one, Cat. No. 780271) using a Hamilton-star liquid handler (Hamilton, Reno, NV, USA). Antibody productivity by the clonal cell lines was assessed in 96 deep-well plates (Greiner bio-one, Cat. No. 780271) using a 10-day fed batch process during which proprietary feeds and glucose were added every other day. On the day of harvest, cells were counted, and the culture supernatant was used for titre measurement using an Octet QK384 (ForteBio, Menlo Park, CA, USA) with Protein A biosensors. The binding rates measured by the Octet were converted to protein concentrations using a standard curve. High and low IgG-expressing CHO clonal populations were identified from the preliminary titre screen (data not shown). Selected clones were scaled up from 96-deep well plates and into 24 square well, V-bottom 10 ml/well polypropylene plates (Porvair Sciences, Cat. No. 360080) and then into Erlenmeyer flasks with vented caps (Corning, Cat. No. 431143)) using proprietary media before being inoculated into a shake flask fed-batch.

### Fed-batch Culture

Antibody production was evaluated from stable CHO pools and clonal cell lines over an 11-day fed-batch regime by seeding cells at 0.7×10^6^ cells/ml in AstraZeneca proprietary medium in a final volume of 45 ml. Cells were cultured at 36.5°C in a humidified 6 % CO_2_ atmosphere in a shaking incubator at 140 rpm. Cell density and viability were monitored on a Vi-Cell XR automated cell counter (Beckman Coulter, High Wycombe, UK). The culture medium was supplemented with bolus additions of a proprietary nutrient feed over the course of the fed-batch process with lactate and glucose monitored using a YSI (2900D, YSI Inc). On day 7, 9 and 11, clarification of 0.5 ml of the cell culture supernatant was performed by centrifugation, and antibody titers were quantified by protein-A HPLC affinity chromatography on an Agilent 1260 Infinity series (Agilent Technologies). To quantify antibody production, the peak size from each sample was compared with a calibration curve. Cell specific productivity (qP) was calculated as follows: QP = Th / CCTf (where Th is the Harvest Titer and CCTf is calculated Cumulative Cell Time on the last day of the culture). CCTi = ((di – di-1) X (VCNi + VCNi-1)/2) + CCTi-1 (where d is the day of the culture, VCN is the viable cell count and i is the day of VCN sampling during the culture).

### Gene Copy Number Analysis

Genomic DNA was extracted from stable CHO pools and clonal cell lines using the PureLink ^TM^ Genomic DNA kit (Invitrogen, USA) according to the manufacturer’s protocol. Gene copy number analysis was carried out on Not1-digested genomic DNA from each sample using the TaqMan Assay System with probes designed against antibody encoding DNA sequence (Thermo Fisher Scientific, USA) and the QX100 Droplet Digital PCR System, along with QuantaSoft software (BioRad). Manufacturer’s instructions were followed throughout.

### Flow Cytometry

Stable pools and clonal cell lines were collected by centrifugation, fixed in 70 % methanol, and stored at -20 °C before further processing. CHO host cells were used as unstained and stained controls. For HC and LC staining, cells were washed twice on ice with DPBS containing 1% BSA, followed by a centrifugation step at 130 x g for 5 min to remove the washing solution. For staining, 2mL 1% BSA prepared in DPBS supplemented with 5 µg/ml APC-conjugated Goat anti-Human IgG Fcγ (Jackson ImmunoResearch, Cat. No. 109-136-170) and 3 µg/ml Kappa-FITC Goat anti-Human antibody (Southern Biotechnology, Cat. No. 2060-02) were added to each sample. Samples were then incubated in the dark on ice for 30-60 min before they were washed twice with 1% BSA. Finally, cell pellets were resuspended in 1 mL of 1 % BSA in DPBS and 200 μL of each sample were transferred into a Nunc™ 96-Well polypropylene microplate (ThermoFisher Scientific, USA,Cat. No. 249944). The stained cells were analysed by a MACSQuant flow cytometer with APC and FITC double-positive population measured according to manufacturer’s specifications. Data analysis was performed using FlowJo software.

### Purification and Aggregate Analysis

Purification of the target protein from cell culture supernatants was performed using PhyTip 200 µl columns, containing 20 µl of ProPlus (MabSelect SuRe™) affinity resin (Biotage GB Limited, Hengoed, UK) operated on a Tecan Freedom EVO® 200 robotic liquid handling platform (TECAN Group Ltd) with Freedom EVOware®, Version 2.7 (TECAN Group Ltd). Column equilibration and first wash used Gibco™ 1x DPBS buffer (Thermo Fisher Scientific). The second wash utilised a buffer composed of 25 mM sodium acetate, 120 mM sodium chloride, pH 5.5. Elution was achieved using 100 mM glycine buffer at pH 2.6. The samples were then neutralised to pH 6.0 with 1M Tris buffer pH 8.0 (Thermo Fisher Scientific). Following purification, aggregation analysis by size exclusion chromatography (SEC) was performed using an ACUITY UPLC SEC 200 column (Waters, MA) on an Agilent 1290 HPLC system. The SEC mobile phase was composed of 50 mM sodium phosphate, 450 mM arginine pH 7.0. Flow rate was set to 0.5 ml/min and injection volume was adjusted to load 40 µg on the column.

### RNA Sample Preparation and Transcriptomic Sequencing

Samples for transcriptomic analysis were taken on day 0 and 6 of the 11-day fed-batch process. Aliquots of 10×10^6^ cells were collected at 130 g for 5 min, the supernatant removed, and the pellet resuspended in 200 µl RNAlater® (Sigma-Aldrich, Cat. No. R0901), snap frozen on dry ice and stored at −80°C before further processing. Upon removal of RNAlater, cell pellets were treated with lysis buffer mix containing Proteinase K sourced from the RNAdvance Tissue kit (Beckman Coulter). Cells were mixed by vortexing, and subsequently incubated at 37°C for 25 min. RNA extraction was performed according to manufacturer’s protocol on a Biomek i7 Hybrid robotic workstation (Beckman Coulter). Subsequent steps involved mRNA enrichment and sequencing library preparation utilizing the KAPA mRNA HyperPrep Kit (Roche), according to the manufacturer’s instructions, using a Tecan Fluent® liquid handler. Library integrity and quality were evaluated using the SS NGS fragment kit (1–6000 bp; Agilent) on a fragment analyzer. Sequencing was performed as paired-end 2×150 reads on an Illumina NovaSeq6000 platform, using a 300-cycle S1 Reagent Kit v1.5 (Illumina).

### RNA-seq Data Processing

For cell pools RNA-seq processing, Fastp (Chen et al., 2018), version 0.22.0, with parameters "--compression 9 --detect_adapter_for_pe--qualified_quality_phred 15 -- unqualified_percent_limit 10 --average_qual 20 --length_required 30", was used to trim PCR and sequencing adapters and filter low quality read pairs. Trimmed and qc filtered reads have then been used by Salmon (Patro et al., 2017), version 1.5.2, with the selective alignment strategy (Srivastava et al., 2020) and the parameters "--libType A --validateMappings – mimicBT2 –useEM", to quantify the expression of transcripts, annotated in Ensembl, version 93 (Howe et al., 2020), with the addition of the extra transgenes. The Bioconductor package txtimport (Soneson et al., 2015) has been used to summarise transcript expression into gene expression. For clonal cell RNA-seq processing, the FASTQ files generated by the sequencer were passed to the nf-core (Ewels et al., 2020) bioinformatics pipeline “rnaseq”, for mapping and quality control. Specifically, version 3.12.0 of the nf-core/rnaseq pipeline was executed, using the 23.04.3 version of Nextflow (Di Tommaso et al., 2017) on a compute cluster running the Slurm scheduler and Singularity containers to host the pipeline software. Sequenced reads were mapped against the Chinese hamster (Cricetulus griseus) reference genome CHOK1GS_HDv1. To quantify expression of the human transgenes, the FASTA sequences of the antibody chains were added to the Chinese hamster reference genome. STAR aligner (Dobin et al., 2012) index files were built from these concatenated genomes, thereby generating a bespoke reference genome for each cell line. For cell pool transcriptomics analysis, raw counts were corrected for batch effect using combat (Zhang et al., 2020) in R before being analysed by DESeq2. For clonal cell transcriptomics analysis, the raw (not normalised) gene-level expression data matrix file generated by the nf-core/rnaseq pipeline was passed to DESeq2 (Love et al., 2014) to identify differentially expressed genes. DEseq2 (version 1.42.1) was used with R (version 4.3.3) on a Debian GNU/Linux 12 (bookworm) operating system, hosted inside a Singularity container. Differentially expressed genes were defined as having changed at least 1.5-fold between the samples being compared, with an adjusted p-value of less than or equal to 0.05.

### Protein Sample Preparation, TMT Labelling and Proteomics Analysis

Cell pellets (10×10^6^ cells) were lysed and homogenised in 0.1 % RapiGest (Waters, USA) buffer as reconstituted in HEPEs buffer (20 mM HEPEs pH 8.0, 2 mM MgCl_2_) supplied with PhoSTOP™ tablets according to manufacturer’s specifications on ice for 30 min. Protein concentration was quantified using a BCA assay (Thermo Fisher, USA). Each sample was diluted to 6 mg/ml in 0.1% RapiGest buffer. To correct batch effects at the end of data analysis, 5 µl from each sample was pooled before aliquoting into 4 tubes each containing 300 µg of protein. 300 µg of internal control or test samples were treated with 500 units Benzonase® Nuclease solution (E1014, Sigma-Aldrich) and incubated at room temperature for 5 min before heat inactivation at 80°C. 200 µg of protein was reduced with 5 mM DTT at 56°C for 30 min and alkylated with 10 mM 2-chloroacetamide (CAA) at room temperature for 30 min. Excess CAA was quenched with 5 mM DTT (final concentration). Lys-C (Promega) was added at a ratio of enzyme:protein of 1:70 (w/w), and incubated for 4 hr at 37°C. Trypsin (Promega) was then added at the same enzyme:protein ratio and digestion was performed overnight at 37°C. Digestion was stopped by the addition of trifluoroacetic acid (TFA) to a final concentration of 0.5 % (pH 2-3) and RapiGest was degraded by heating at 37°C for 45 min. Precipitates were removed by centrifugation at 18000x g for 8 min. Supernatants of cell lysates were desalted using home-made C18 stage tips (3M Empore) packed with poros R3 resin (Thermo Scientific). Stage tips were equilibrated with 80 % acetonitrile (MeCN)/0.5 % FA followed by 0.5 % FA. Bound peptides were eluted with 30-80 % MeCN/0.5 % FA and lyophilized.

### Tandem Mass Tag (TMT) Labeling

Dried peptide mixtures from each condition were resuspended in 90 µl of 200 mM HEPEs, pH 8.5. 45 µl TMTpro 18plex reagent (Thermo Fisher Scientific,UK), reconstituted according to manufacturer’s instructions, was added to each sample and incubated at room temperature for an hour. The labeling reaction was then terminated by incubation with 9 µl 5 % hydroxylamine for 30 min. The labeled peptides were combined into a single TMT multiplex and were desalted using the same stage tips method as above.

### Off-line High pH Reverse-phase Peptide Fractionation

Experiments were carried out using XBridge BEH130 C18, 5 µm, 2.1 x 150 mM (Waters) column with XBridge BEH C18, 5 µm Van Guard cartridge, connected to an UltiMate 3000 Nano/Capillary LC System (Dionex). 100 µg of the labeled peptides were reconstituted in 5 % MeCN/10 mM ammonium bicarbonate and were separated with a gradient of 3-90 % B (A: 5 % MeCN/10 mM ammonium bicarbonate, pH 8.0; B: MeCN/10 mM ammonium bicarbonate, pH 8, [9:1]) for 60 min at a flow rate of 200 µl/min. Fractions were collected every minute and combined into a total of 18 fractions, then lyophilised. Dried fractionated peptides were resuspended in 1 % MeCN/0.5 % FA, desalted using C18 stage tips before mass spectrometry analysis.

### Mass Spectrometry Analysis

High pH reverse-phase fractionated peptides were analysed by LC-MS/MS using a fully automated Ultimate 3000 RSLC nano System, fitted with a PepMap Neo C18, 5 μm 0.3 x 5 mm nano trap column (Thermo Fisher Scientific,UK) and an Aurora ultimate TS 75 μm x 25 cm x 1.7 μm C18 column (IonOpticks). Peptides were separated using buffer A (0.1 % FA) and buffer B (80 % MeCN, 0.1 % FA) at a flow rate of 300 nl/min and column temperature of 40°C. Eluted peptides were introduced directly via a nanoFlex ion source into an Orbitrap Eclipse mass spectrometer (Thermo Fisher Scientific,UK). The mass spectrometer was operated in real-time database search (RTS) with synchronous-precursor selection (SPS)-MS3 analysis for reporter ion quantification. MS1 spectra were acquired using the following settings: Resolution=120K; mass range=400-1400m/z; AGC target=4e5; MaxIT=50ms and dynamic exclusion was set at 60s. MS2 analysis were carried out with HCD activation, ion trap detection, AGC=1e4; MaxIT=50ms; NCE=33% and isolation window =0.7m/z. RTS of MS2 spectrum was set up to search UP000001075 Chinese Hamster proteome (08 Feb 2023), with fixed modifications cysteine carbamidomethylation and TMTpro 16plex at N-terminal and Lysine residues. Methionine-oxidation was set as variable modification. Missed cleavage=1 and maximum variable modifications=2. MS3 scans were performed with the close-out function enabled and max number of peptides per protein = 5. The selected precursors were fragmented by HCD and analyze using the orbitrap with these settings: Isolation window=0.7 m/z; NCE=55, orbitrap resolution=120K; scan range=110-450 m/z; MaxIT=300ms and AGC=1.5 e^5^.

### MaxQuant Data Processing

The raw LC-MS/MS data were processed using MaxQuant (Cox & Mann, 2008) with the integrated Andromeda search engine (v.2.4.2.0) in house at Medical Research Council Laboratory of Molecular Biology. MS/MS spectra were quantified with reporter ion MS3 for proteomics from TMTpro 18plex experiments and searched against UP000001075_Chinese Hamster FASTA database (downloaded on 08 Feb 2023). Carbamidomethylation of cysteines was specified as fixed modification, with methionine oxidation and N-terminal acetylation (protein) set as variable modifications. Protein quantification requirements were set at 1 unique and razor peptide. Other parameters in MaxQuant were kept as default values. MaxQuant data was then imported into Perseus (v.2.0.10.0) to remove identifications from reverse database, identifications with modified peptide only (for protein groups) and common contaminants. Data with the reporter intensities of ‘0’ were converted to NAN and exported as text files for further data analysis.

### Omics Analysis

iDEP (2.01) R package analysis tool (Ge et al., 2018) was used to perform PCA, heatmap, k-mean clustering and GO term enrichment analysis for both RNA-seq and proteomics data. Briefly, when analysing RNA-seq data from either cell pools or clones, batch effect corrected raw counts were VST transformed before PCA and k-mean clustering were performed. When analysing proteomics data, background corrected and normalized abundances were Log2 transformed with missing values replaced by group median before PCA and k-mean clustering were performed. k-mean clustering analysis was carried out by choosing the number of top variable genes or proteins whose standard deviation were larger than the background SD value followed by Elbow’s method to identify the minimum number of clusters that can summarise the variation from the top variable proteins or genes. The heatmap was plotted based on VST-transformed or log2 transformed data being Z-scored i.e. by dividing row mean with SD. Genes or proteins from each cluster were interrogated for enrichment of GO terms by using iDEP or ShinyGO (0.80) (Ge et al., 2020) R package analysis tools with list of filtered genes and proteins used as background.

For differential gene or protein expression analysis, DESeq2 package was applied to RNA-seq data for both cell pools and clones as described above. For differential protein expression analysis of cell pools, Proteome Discoverer (PD) (Thermo Fisher, USA) normalized protein abundances were transformed using edgeR: Log2(abundance+4) before pair-wised analysis was carried out using limma with voom. For differential protein expression analysis of clones, proteomic abundances from MaxQuant were input into the proteomics analysis software Lava (H.T. Parsons ; https://github.com/tempeparsons/Lava). Lava is a pair-wise test algorithm that performs 2-sample test on Z-score normalized Log2 data with batch effect corrected by following a ratio based scaling method(Yu et al., 2023). Any missing values remained at zero. When comparing two groups of (corrected and normalised) replicate abundance observations for each protein/peptide, p-values were calculated for all groups, including those with a few missing values, by employing a t-test that tolerates unequal group sizes using the SciPy function ’ttest_ind_from_stats’ (https://docs.scipy.org/doc/scipy/reference/generated/scipy.stats.ttest_ind_from_stats.html). Proteins detected in at least one out of three replicates of each group with peptide counts >2 was selected and plotted in volcano plot.

### Data Visualization

Volcano plots generated by ggplot2 in R (4.3.1) were used to visualize differentially expressed genes or proteins. Enrichment of GO terms were assessed by importing DE genes or proteins into ShinyGO (0.80) (Ge et al., 2020). Venn diagrams were plotted using VennDiagram package in R (4.3.1).

### Statistical Analysis

Statistical analyses were performed using GraphPad Prism (GraphPad, USA), version 10.2.3. Results are expressed as meanC±Cstandard error of the mean (SEM). One-way ANOVA was used for qP and high molecular weight species comparisons with Tukey’s post-hoc multiple comparison method applied. The significance levels between cell pools are indicated by asterisks in each figure above the bars: *pC<C.05; **pC<C.01; ***pC<C.001. Benjamini and Hochberg’s method was used to generate FDR for both transcriptomics and proteomics analysis, and significance level for BiP expression was represented by FDR with ** p <.01, ***pC<C.001, ****p <.0001. Correlation analysis of clonal cell line gene copy number, mRNA and protein abundances were calculated using the Pearson correlation coefficient method.

## Results

### Generation of Cell Lines Expressing ETE and DTE Biotherapeutics

To investigate cellular adaptations that occur in response to the expression of ETE and DTE biotherapeutics, we generated a panel of monoclonal and bispecific antibodies with differential expression capabilities. These included one known ETE monoclonal antibody (mAb1), two known DTE bispecific antibodies (BisAb1 and BisAb2), and a monovalent control for each of the bispecific antibodies (MonoAb1 and MonoAb2, respectively) whose expression levels were unknown (**Figure 1A**). Stable, random DNA integration pools expressing each of the five molecules were generated and assessed in fed-batch culture to measure cell growth and antibody expression, and to generate material for multi-omics characterisation according to the schematic shown (**Figure 1B**). Stable pools in fed-batch revealed different performance data across different antibody producing cell lines. Cells expressing mAb1 reached peak viable cell density (VCD) on day 7, compared to other cell pools that reached peak VCD on day 9. BisAb1- and MonoAb2-expressing cell pools displayed relatively low growth rates compared to other cell pools. Consistent viability profiles were observed across all pools apart from mAb1-expressing cells where viability sharply declined after day 9 (**Figure 1C**). Assessment of intracellular HC and LC levels by flow cytometry demonstrated homogenous co-expression of both heavy and light chains in all pools apart from those expressing BisAb2 where multiple cell populations were observed (**Supplementary Figure 1**). Measurement of secreted antibody titre revealed that mAb1, a known ETE antibody, had the highest peak titre of 2069mg/L. By contrast all other cell pools produced significantly lower peak titres ranging from 22mg/L to 868mg/L, with BisAb2-expressing pools yielding the lowest titre (**Figure 1D**). To assess antibody secretion according to the amount of antibody produced per cell per day, cell-specific productivity (qP) was calculated. Here, consistent with the data for volumetric titre analysis, mAb1- and BisAb2-expressing pools displayed the highest and lowest qP respectively (**Figure 1E**). Interestingly, the low qP of BisAb1 and BisAb2 was partially rescued when they were converted to their monoclonal antibody controls (MonoAb1 and MonoAb2, respectively) (**Figure 1E**). Finally, the secreted antibody from each cell pool was purified and examined for the presence of higher molecular weight species (aggregates) to give an indication of product quality. Each molecule was compared to the mAb1 ETE control. MAb1 showed no aggregates, followed by MonoAb1 with aggregation level of 7.4% ± 0.65. By contrast, BisAb1 displayed aggregation values at 28.5% ± 0.01, and MonoAb2 and BisAb2 displayed the highest levels with values of 51.4% ± 1.64 and 51.6% ± 7.72 respectively (**Figure 1F**).

### Multi-omics Assessment of Cell Pools Revealed a Limited Number of Signatures Associated with ETE and DTE Antibody Expressing Cell Lines

To discover biological signatures that were associated with ETE and DTE phenotypes, we collected multi-omics data, encompassing transcriptomic and proteomic analyses, of all cell pools. Because BiP is a known chaperone for IgG biosynthesis, and strongly influences protein folding in addition to being a primary target of the ER-stress induced unfolded protein response (UPR), we first examined BiP transcript and protein levels across the cell pools (**Figure 2A**). Antibody expressing cell lines showed a small but significant elevation in BiP transcript and protein levels compared to no IgG controls. Seeking to further elucidate differences between antibody-expressing and non-expressing pools, we performed a pair-wise comparison of each expressing cell pool to the no IgG control pool. Both transcriptomic and proteomic analyses resulted in a limited number of significantly differentially expressed genes or proteins that could be used for pathway analysis (**Supplementary Figure 2A**). At a threshold of false discovery rate (FDR) <0.05 and fold change of >1.2, each cell pool displayed a small number of differentially expressed genes and proteins, with transcriptomics returning more differentially expressed signatures than proteomics. These signatures were associated with processes such as oxidative phosphorylation and protein folding in the ER (**Supplementary Figure 2B**). To examine signatures associated with ETE and DTE outcomes, pools were grouped according to expression levels of the biotherapeutic, with cell pools displaying significantly lower qPs than mAb1 being defined as DTE and all others as ETE (**Figure 1D**). Principal component analysis (PCA) was performed on transcriptomic and proteomic data sets (**Figure 2B**) but revealed no clear separation of ETE and DTE lines. We then reviewed the transcriptomic data for genes that were differentially regulated across ETE or DTE cell pools as compared to non-expressing control cells (**Figure 2C, Supplementary Figure 2C**). Here, Mt_rRNA, ND6, ND3 and Gas7 represent upregulated genes and Wsb2 and Col6a1 represent downregulated genes shared by at least two ETE cell lines. Biological process was explored for each gene where ND3 and ND6 represent mitochondrial genes that are involved in electron transport whereas Gas7 is involved in differentiation of cerebellar neurons and was previously shown to be upregulated in a mitochondrial enriched CHO host (Chakrabarti et al., 2022). Wsb2 is involved in ubiquitin conjugation pathways whereas Col6a1, an extracellular matrix protein, is involved in cell adhesion. By contrast, Mt_rRNA, Fscn1 and Synpo2 represent upregulated genes and C3, Cgnl1, C1qtnf1, Itga3 and Cited1 represent downregulated genes shared by DTE cell lines. Fscn1 and Synpo2 are involved in actin filament bundle formation and assembly and are important for cell migration and focal adhesion. C3 is involved in complementary pathway and fatty acid metabolism and was shown to be downregulated in CHO subcloning processes (Weinguny et al., 2020). Cited1 is involved in apoptosis and cell differentiation, Itga3 functions as a cell receptor that is involved in cell adhesion, C1qtnf1 is involved in negative regulation of platelet aggregation (Lasser et al., 2006) and a related family member was reported to be downregulated in later growth phases in CHO clones (Lin et al., 2020). Cgnl1 is involved in anchoring the apical junctional complex to actin-based cytoskeletons. The small number of significantly altered genes/proteins shared by either ETE or DTE expressing pools leads us to conclude that each antibody expressing pool, derived from random integration of the antibody expressing transgene is highly heterogeneous and that this might mask discovering cellular adaptations which could act on a cell-by-cell basis.

**Figure 2:**
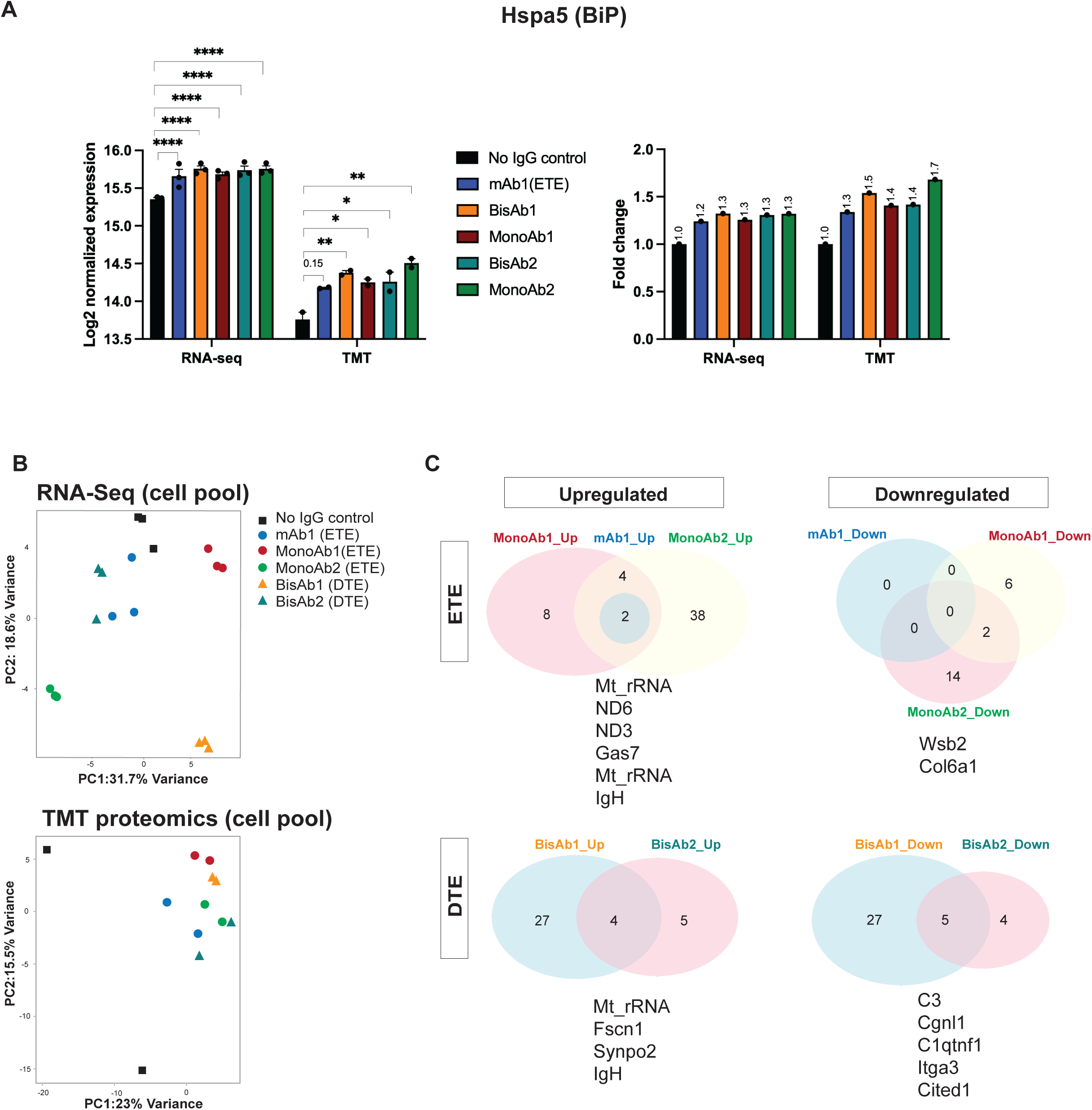
RNA-seq and TMT Proteomics Analysis of Cell Pools. **A:** Comparison of Hspa5 (BiP) mRNA (RNA-seq) and protein expression (TMT) of each cell pool with No IgG expressing control pools, data presented as Log2 normalised expression and fold change. **B:** PCA analysis of cell pools from RNA-Seq and TMT proteomics. **C:** Venn diagram derived from RNA-seq data detailing the number and names of differentially expressed genes shared by at least two ETE and two DTE biotherapeutic expressing cell lines.

### Generation of Clonal Cell Lines and Selection for High and Low Secretion Phenotypes

To overcome the under-representation of robust cellular signatures within our pooled lines, we generated clonal cell lines expressing each of our target biotherapeutics and assessed a panel of clones for antibody productivity and product quality (**Figure 3A**). Clonal cell lines were evaluated in fed-batch to identify high and low qP clones with comparable growth profiles and intracellular HC and LC content (**Supplementary Figure 3 and 4**). High and low qP clones were identified for all lines apart from BisAb2 where no high secreting clones were found (**Figure 3B**). Assessment of product aggregation revealed that clonal cell lines expressing bispecific antibodies showed high levels of extracellular aggregates compared to clonal cell lines expressing monoclonal antibodies where negligible extracellular aggregates were observed. BisAb1 and BisAb2 expressing clones led to 30.62% ± 2.41 and 11.45% ± 0.29 extracellular aggregates respectively (**Figure 3C**). Interestingly, MonoAb2 expressing clones led to 1.81% ± 0.28 extracellular aggregates, a result not observed in the corresponding pools where average aggregation was 51.62% (**Figure 1F**).

**Figure 3:**
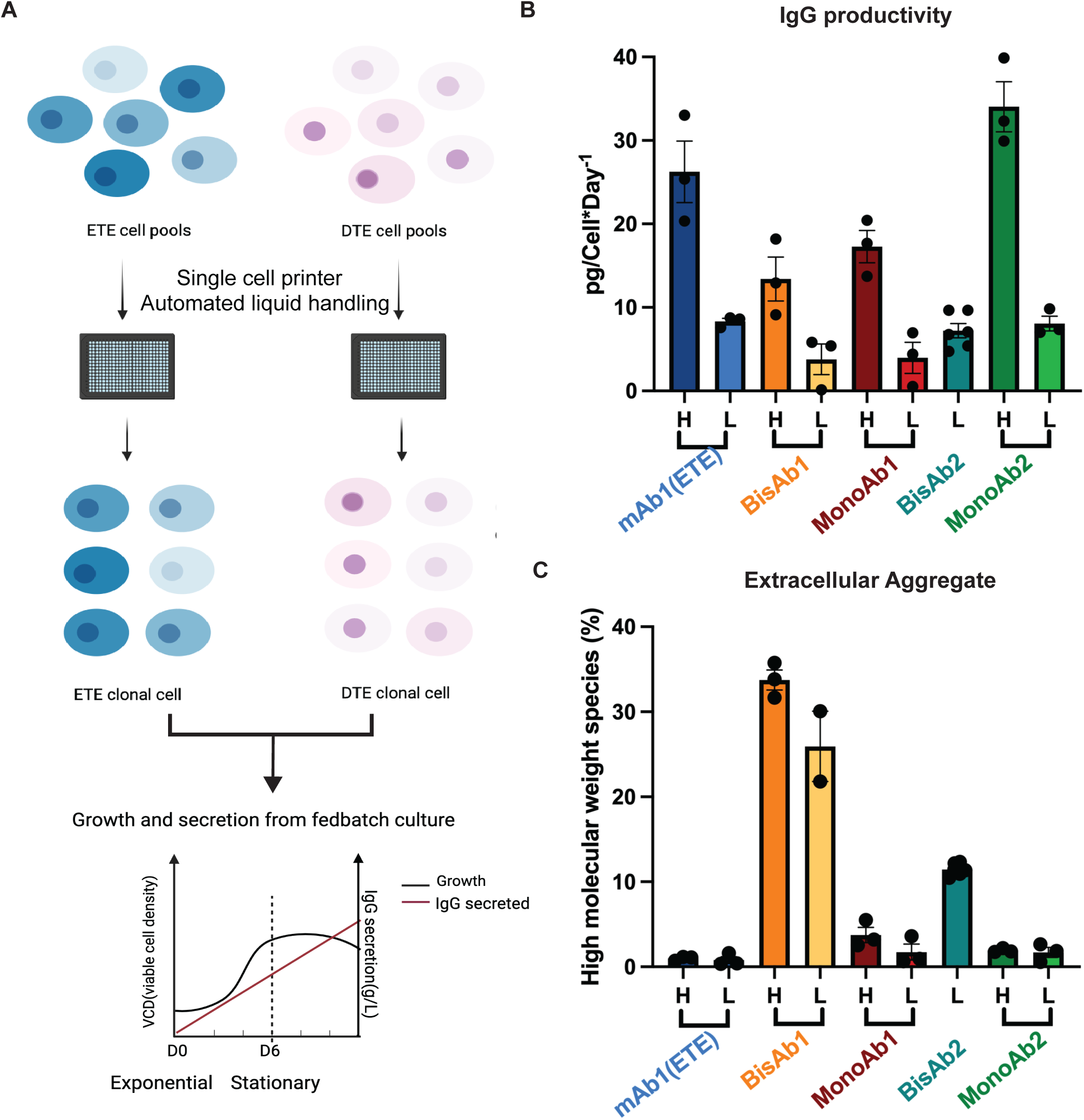
Clonal Cell Line Generation and Characterisation. **A:** A schematic of the clonal cell line generation process detailing the selection of clones with various secretion capabilities for downstream characterisation studies. **B:** High (H) and low (L) antibody secreting clones for each biotherapeutic were grouped according to cell specific productivity (qP) data derived from a fed-batch culture evaluation. **C:** Biotherapeutic high molecular weight species percentage derived from each clonal cell line. Data presented as mean ± standard deviation (SD) from N=3 or N=2 clone from either high qP or low qP group of each cell line.

### Multi-omics Analysis of Clonal Cell Lines

To discover biological signatures associated with clone secretion phenotypes, proteomic analysis was performed on lysates from D0 and D6 of fed-batch culture. Initial unbiased k-mean clustering analysis of proteomics data showed that the culture time was the largest contributor to variation among samples (**Figure 4A**). Here, when cells reached day 6 in culture, proteins with functions in lipid catabolic processes as well as carbohydrate derivative catabolic processes were upregulated while proteins with functions in transcription, DNA-templated regulation of macromolecule biosynthetic processes as well as cell cycle were downregulated. Given the metabolic changes that occur during the fed-batch process, it is perhaps not surprising that the dominant signature was time-point of cell collection. We reasoned that this dominant effect may prevent identification of clusters that result from high and low qP phenotypes. To address this, we separated the D0 and D6 samples for independent analysis. For both D0 and D6, clonal lines did not cluster according to qP, with individual clones expressing the same biotherapeutic not necessarily clustering together (**Figure 4B**). Similar lack of clustering of high and low qP clones was observed in the transcriptomics dataset (**Supplementary Figure 5A and B**).

**Figure 4:**
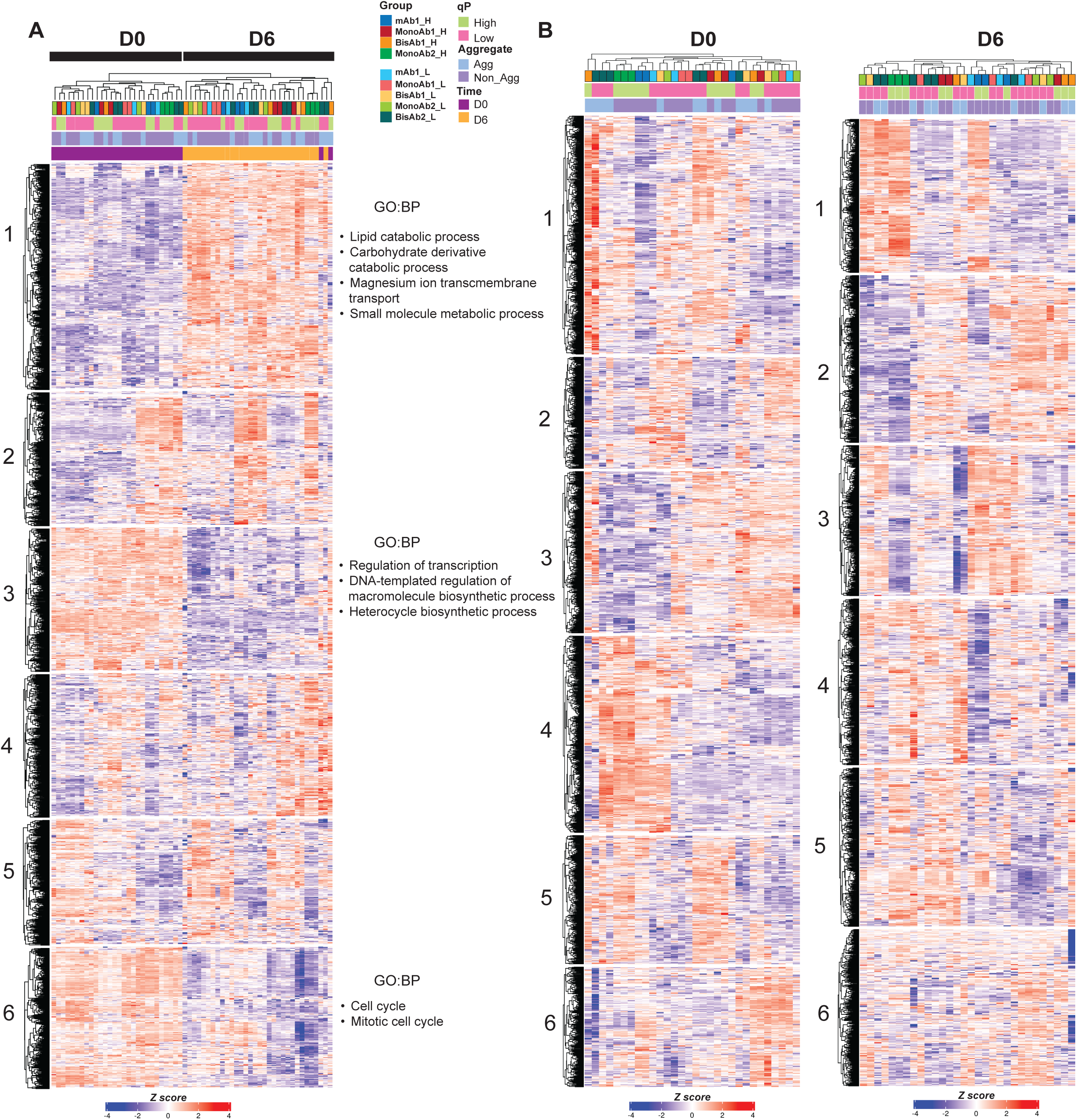
Proteomics Analysis from Clonal Cell Lines. **A-B:** Heatmaps derived from TMT proteomics for clonal cell lines analysed on day 0 (D0) and day 6 (D6) of the fed batch process. Data displayed by combining D0 and D6 (**A**) or displayed independently of day in culture (**B**). K-mean clustering was applied to data, enriched GO terms with biological process (BP) from cluster 1/3/6 are highlighted.

### Gene Copy Number, Transgene mRNA Expression and Protein Abundance are Primary Drivers of qP in Clonal Cell Lines

Since our multi-omics analysis of clones failed to identify relationships among high qP expressing clones, we hypothesised that the high secretion phenotype might be dominated by stochastic properties of different clones. Preliminary observations of the proteomic and RNA-seq datasets suggested HC abundance was related to qP (data not shown). We therefore characterized the clonal lines with respect to transgene copy number, mRNA transcript abundance, and protein expression. A large variation in gene copy number, transcript level and protein abundances were observed across all clones (**Supplementary Figure 6A-C**). As might be expected, mRNA abundance for LC and HC correlated well with gene copy number (GCN): r=0.4717, p<0.001 for LC, and r=0.522, p<0.001 for HC (**Figure 5A-B**). Similarly, we observed a significant positive correlation between mRNA and protein abundance for LC (r=0.439, p<0.00001) (**Figure 5C**). When HC mRNA and protein abundances were compared, correlation was poor, but we noted two divergent correlations associated with cell line (**Figure 5D**), with BisAb2/MonoAb2 separated from the others. We re-calculated correlations separately for these two groups, revealing significant correlation between HC mRNA and protein abundances in a cell line specific manner (r=0.8822, p<0.00001; r=0.9606, p<0.00001) (**Figure 5E-F**). Taken together, these data suggest that transgene expression is a dominant determinant of qP in most clonal cell lines, with higher gene copy number giving rise to higher mRNA levels and higher protein abundance.

**Figure 5:**
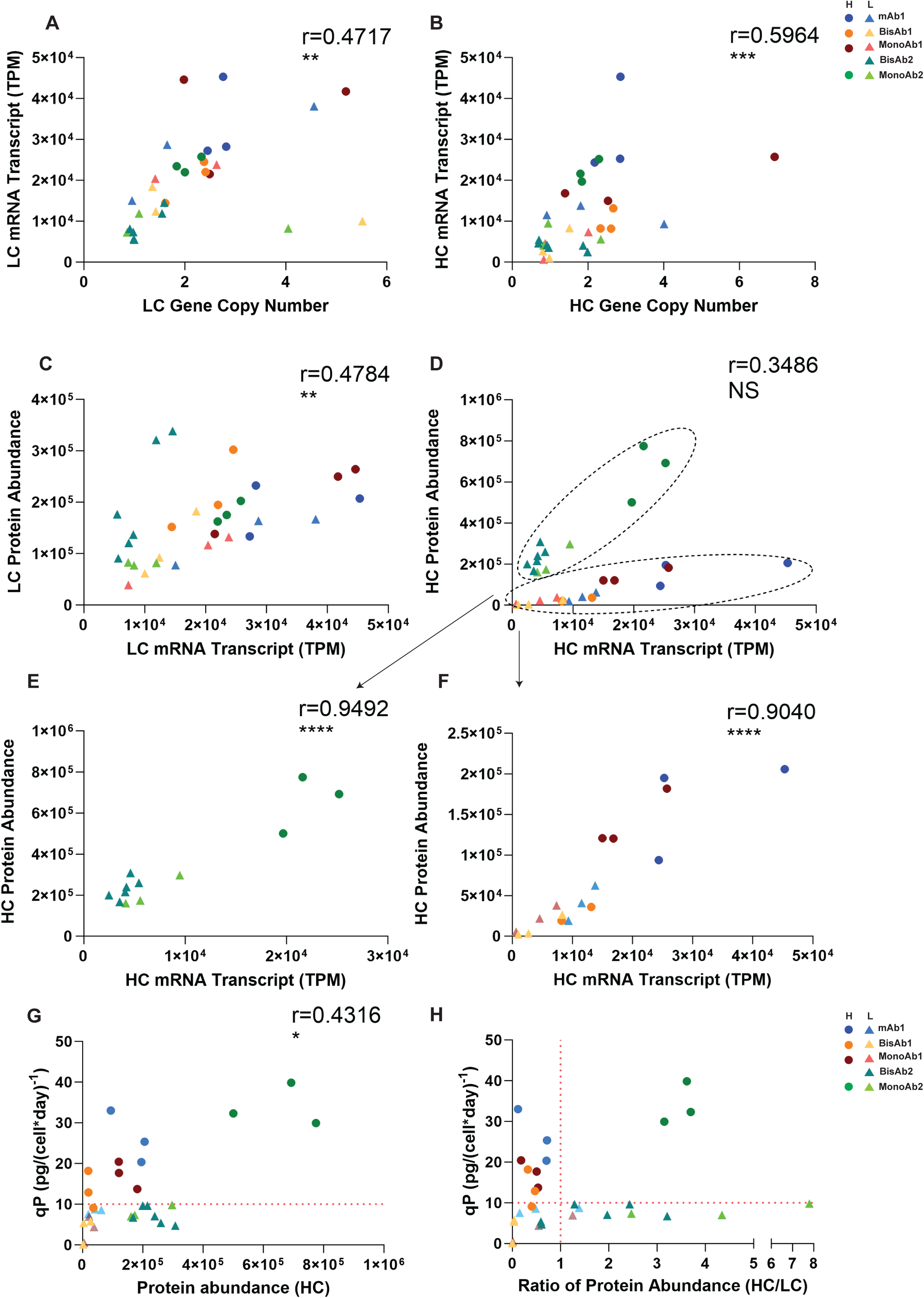
Correlation of Clonal Cell Line Gene Copy Number, mRNA Abundance, Total Protein Abundance and qP. **A-B:** Correlation of gene copy number and mRNA abundance for light chain (**A**) and heavy chain (**B**) for all clones. **C-D:** Correlation of mRNA abundance with protein abundance for light chain (**C**) and heavy chain (**D**) for all clones. **E:** Correlation of mRNA abundance with protein abundance for heavy chain from clones expressing MonoAb2 and BisAb2. **F:** Correlation of mRNA abundance with protein abundance for heavy chain from clones expressing mAb1, MonoAb1 and BisAb1. **G:** Correlation of protein abundance of heavy chain at D0 with the cell specific productivity (qP) from each clonal cell line. **H:** Protein ratio of heavy chain to light chain plotted against the cell specific productivity (qP) from each clonal cell line. Pearson correlation coefficient was used to test significance in correlation. P value *<0.05. **<0.01, *****<0.0001

Lastly, we sought to determine the relationship between protein abundance and secretory capacity. Plotting HC and LC abundance relative to qP, we found no correlation between LC abundance and secretion yield (**Supplementary Figure 6D**), but a reasonable correlation between HC abundance and qP (**Figure 5G**). This suggests that HC abundance is a key component of secretion yield, which is consistent with known behaviour of IgG assembly, where unpaired HC is retained in the ER, thereby limiting secretion. Previous studies showed that an imbalance in heavy and light chain expression is known to impact both yield and aggregation (Eisenhut et al., 2020; Leon P. Pybus et al., 2014; Schlatter et al., 2005). We therefore tested whether the HC/LC ratio changed with qP, expecting to find that an optimal ratio of 1:1 would be associated with higher qP. Instead, we found little relationship between qP and HC/LC ratio and were surprised to see one set of high secretors (MonoAb2) with a high HC/LC ratio, suggesting excellent secretion despite significant overexpression of HC (**Figure 5H**).

### Identification of Biological Signatures Associated with High qP Clones

Having defined several important drivers of HC/LC expression levels, we returned to our proteomic data, comparing high and low secretion clones for each individual biotherapeutic rather than grouping all high expressors together. This pair-wise comparison revealed proteins differentially expressed in high secreting clones for each cell line. Differential protein abundance between high and low secretors expressing the same biotherapeutic was visualised using volcano plots (**Supplementary Figure 7**). In each case, a small number of proteins were significantly different in their abundance: either more abundant in high secretors, or more abundant in low secretors (**Supplementary Figure 7**). Of these several were shared across all cell lines regardless of the biotherapeutic being expressed (**Figure 6A**). Shared factors that were more abundant in high secretors include Foxo3, Fgfr1op2 and Spryd3 at D0, and Clcn4, Tmem104, Arhgap33 at D6 (**Figure 6A, left**). **p a**Pr**n**ot**e**ei**l**n**s**s that were shared as more abundant in low qP clones include Ngrn, Foxm1, Pkp3 at D0, and Tmem39b and Map9 at D6 (**Figure 6A, right panels**).

**Figure 6:**
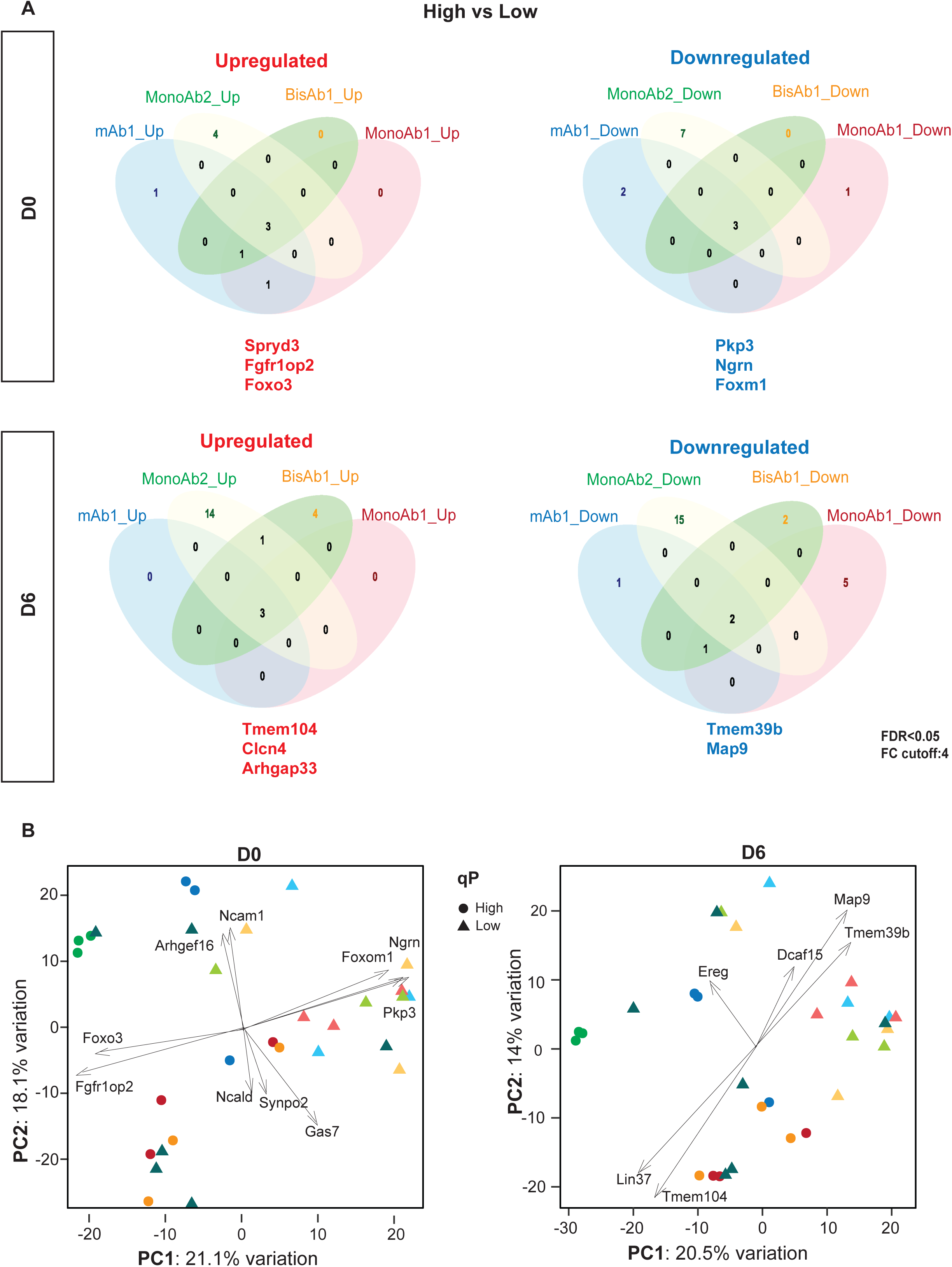
Identification of Protein Hits Associated with High and Low qP. **A**: Venn diagrams derived from D0 and D6 proteomics data detailing significant up and downregulated proteins in high secreting clones that are shared by 4 cell lines: mAb1, MonoAb1, MonoAb2 and BisAb1 as well as the number of proteins specific to each cell line. Significant proteins were defined by having a fold change > 4 and adjusted-P value (FDR) <0.05. **B**: PCA analysis of proteomics data from D0 and D6. Two components with significant correlation with qP (PC1) and cell line (PC2) are shown.

Principal component analysis (PCA) on proteomics data from either D0 or D6 (**Figure 6B**) successfully separated cell lines by PC1 (correlated with qP) and PC2 (correlated with cell line). This analysis identified candidate drivers that might contribute to separation of clones according to the two components shown. Drivers associated with the component representing qP (PC1) included Foxo3 and Fgfr1op2 (correlated with high qP) and Foxm1, Ngrn and Pkp3 (correlated with low qP) at D0 (**Figure 6B, left pan**)**e**. **l**For the D6 analysis, Lin37 and Tmem104 were associated with high qP, and Map9, Tmem39b and Dcaf15 associated with low qP (**Figure 6B, right panel**). The correspondence between select candidate PCA drivers and factors shared by all high or low qP clones suggests that these hits may represent important individual proteins that contribute to secretion phenotypes.

## Discussion

Efforts to enhance recombinant protein production in CHO cells have primarily concentrated on engineering transgenes and vectors (Blanco et al., 2020; Zhong & D’Antona, 2021), and refinement of bioprocess conditions such as pH, temperature and media and feed composition (Gomez et al., 2018; Lee et al., 2021). Although several studies have explored the manipulation of host cellular pathways to boost yields of both monospecific and multispecific antibodies (Budge et al., 2020; Florin et al., 2009; Johari et al., 2015; Orellana et al., 2015; L. P. Pybus et al., 2014), the cellular mechanisms coordinating host cell genetic adaptations to high levels of secretion and varying formats of biotherapeutics are not yet fully understood. Here, aiming to gain further insights into biological signatures associated with high secretion phenotypes, we analysed engineered cell lines that express a panel of ETE and DTE biotherapeutic antibody formats.

Since our initial stable cell pools were too heterogeneous to yield robust biological signatures, we focused on clonal lines, derived from the initial pools and selected for both high and low production phenotypes. Quantitative transcriptomics and proteomics analysis of these clones grown under production conditions showed that culture time was the largest contributor to variation among samples. Both transcript and protein expression profiles clustered according to culture phase: seeding (day 0) and exponential growth (day 6). The metabolic profiling of CHO cells during the production of biotherapeutics in fed batch culture has been extensively characterized (Coulet et al., 2022). Different cell growth phases have distinctive metabolic pathways, reflected in the specific proteins involved in these circuits (Gopalakrishnan et al., 2024). It is therefore not surprising that the dominant signature across all antibody-expressing clones, independent of secretion profiles, was time-point of cell collection. Surprisingly, when this growth-phase dependency was removed, by separating the analyses of D0 and D6 samples, we still failed to observe clear relationships linked to high and low qP clones. The observation that in some cases, individual clones expressing the same biotherapeutic antibody failed to cluster together, prompted us to consider that each clone carries a completely unique epigenetic and genetic complement, with inherent gene expression stochasticity for the observed cell-to-cell variability in these clonal populations revealing a major problem for identifying signatures that might be manipulated for commercial advantage.

Pairwise comparison of the high- and low-secreting clones for each biotherapeutic allowed us to identify several candidate drivers associated with productivity, some of which haven’t previously been reported in studies related to biotherapeutic production in CHO. Forkhead Box O3 (FOXO3), SPRY domain containing 3 (SPRYD3), FGFR1OP2, Clcn4, Arhgap33, and Tmem104 were all more abundant in high qP clones. How these proteins impact secretion remains to be determined and they do not seem to be involved in a shared pathway. However, the individual functions of these proteins might be revealing. FOXO3 is a transcription factor that promotes autophagy (Mammucari et al., 2007; Zhao et al., 2007; Zhou et al., 2012), which might promote degradation of misfolded internal aggregates, thereby enhancing secretion of correctly assembled antibodies. Indeed, FOXO signalling was previously reported to be associated with qP (Tamošaitis & Smales, 2018) and a recent study described the addition of autophagy-inducing peptides (Braasch et al., 2021) to enhance productivity. Clcn4 has been described as a Cl^−^/H^+^ exchanger that predominantly resides in the ER and is involved in endosome trafficking (Guzman et al., 2017). Tmem104 is another ER membrane protein whose biological function remains unknown. Finally, Arhgap33 has recently been described as a scaffolding protein that facilitates the formation of Cdc42/SORT1-containing trafficking complexes to enhance selective protein transport from the Golgi apparatus (Nakazawa et al., 2016), implicating post-Golgi transport as another avenue for potential intervention.

Proteins overexpressed in low-secreting clones included Foxm1, Ngrn, Pkp3, Map9 and Tmem39B. Foxm1 is a transcription factor expressed in actively dividing cells and is critical for cellular functions such as proliferation and cell cycle progression (Kalin et al., 2011). Ngrn has been associated with mitochondrial ribosome biogenesis, specifically regulating the mitochondrial 16S rRNA and intra-mitochondrial translation (Arroyo et al., 2016). Map9 has been associated with chromosomal instability (Wang et al., 2020), whereas Tmem39b (Zhuang et al., 2024) has been described as a cell proliferation regulator. In line with our observations, previous studies demonstrated that modulating cell cycle (Kumar et al., 2007), cell proliferation (Kaufmann et al., 2001; Khoo & Al-Rubeai, 2009; Meents et al., 2002), chromosome instability (Huhn et al., 2022), autophagy (Barzadd et al., 2022; Braasch et al., 2021), and mitochondrial activity (Chakrabarti et al., 2022; Chakrabarti et al., 2019; Dhiman et al., 2019), can help improve the expression of a recombinant proteins in CHO cells.

Overall, we conclude that heterogeneity even among related clones represents a confounding effect that obscures biological signatures associated with high productivity. This agrees with previous studies demonstrating that subclones undergo diversified adaptations resulting from a combination of genomic, transcriptomic as well as epigenetic changes (Marx et al., 2022; Weinguny et al., 2020). Despite this, novel driver proteins were identified that could represent future engineering targets to help modulate processes such as post-Golgi trafficking, autophagy, cell proliferation and mitochondrial ribosome biogenesis and may therefore help fine-tune the productivity of novel clonal lines.

## Author contributions

R.K.M., E.A.M., Y.B., and I.D.M. conceptualized and designed the experiments; Y.B. and I.D.M. performed cell line generation and phenotypic characterisation experiments; Y.B. performed protein sample preparation before TMT labelling, S.Y.P and C.F performed TMT labelling, mass spectrometry and proteomic analysis; S.W., T.S., Y.B. and L.G. contributed to omics data analysis, data visualization and statistical analysis; G.L. contributed to aggregate quantitation from cell culture media; R.E. conducted RNA sample preparation and sequencing library preparation for transcriptomic analysis; N.H. helped with ddPCR sample preparation for Gene copy number analysis and clonal cell line generation; C.S.F. generated the mAb1 CHO stable cell line; Y.B., I.D.M., R.K.M., and E.A.M. wrote and reviewed the manuscript. D.H reviewed the manuscript; R.K.M., and E.A.M. conceived the project.

## Acknowledgments

The authors thank Fabio Zurlo and the Bioprocess assay team at AstraZeneca for performing HPLC titer analysis; Dr Hattie Parsons for developing the Lava pipeline for proteomics analysis; all the members from Liz Miller’s group at the MRC Laboratory of Molecular Biology for very insightful scientific feedback; all the members of the Cell Line Development team at AstraZeneca for support and helpful discussion; and Kerensa Klottrup-Rees for providing guidance and support in the generation of clonal cell lines.

## Data and materials availability

All data are available in the main text or the supplementary materials.

---

**Supplementary Figure 1: Intracellular IgG Expression in Cell Pools.** Intracellular expression of heavy chain (HC) and light chain (LC) for No IgG control, mAb1, BisAb1, MonoAb1, BisAb2 and MonoAb2 pools as determined by flow cytometry. Quadrant 1 (Q1) corresponds to cells with only kappa LC expression, quadrant 2 (Q2) corresponds to double positive cells expressing both HC and kappa LC, quadrant 3 (Q3) corresponds to cells expressing only HC, quadrant 4 (Q4) corresponds to non-expressing cells. Q4 was determined based on naïve CHO cells stained with only secondary antibodies and unstained naïve CHO cells.

**Supplementary Figure 2: Cell Pool Transcriptomic and Proteomic Analysis. A:** A bar chart indicating the number of significantly differentially expressed proteins and genes in cell pools as measured from TMT proteomics or RNA-seq. **B:** A heatmap showing significantly upregulated genes shared by all antibody producing cell lines when compared to the no IgG control with significantly expressed genes defined as adjusted-P value <0.05 and a fold change > 1.2. **C:** Volcano plot of differentially expressed genes from each cell line compared to No IgG control. Adjusted-P value (FDR) <0.05 and fold change > 1.5 were used to define significantly expressed genes. Upregulated genes from each antibody producing cell line were highlighted in red while downregulated genes from each antibody producing cell line were highlighted in blue. Log2FC and - Log10 (P-adjusted value) were used to construct volcano plots.

**Supplementary Figure 3: Clonal Cell Line Characterisation Data used for Selection of Clones for Omics Analysis.** Clonal cell line titre, growth (viable cell density, VCD), viability and cell specific productivity (qP) derived from day 7, 9 and 11 of the fed batch process for mAb1 (**A**), BisAb1 (**B**), MonoAb1 (**C**), BisAb2 (**D**), MonoAb2 (**E**). Asterisk displayed on qP graphs denote clonal lines used for omics analysis with high qP clones in red and low qP clones in blue.

**Supplementary Figure 4: Intracellular IgG Expression in Clonal Cell Lines.** Intracellular expression of heavy chain (HC) and light chain (LC) for mAb1, BisAb1, MonoAb1, BisAb2 and MonoAb2 expressing clonal cell lines as determined by flow cytometry. Quadrant 1 (Q1) corresponds to cells with only kappa LC expression, quadrant 2 (Q2) corresponds to double positive cells expressing both HC and kappa LC, quadrant 3 (Q3) corresponds to cells expressing only HC, quadrant 4 (Q4) corresponds non-expressing cells.

**Supplementary Figure 5: Transcriptomic Analysis of Clonal Cell Lines. A-B:** Heatmaps derived from RNA-seq data from clonal cell lines analysed on day 0 (D0) and day 6 (D6) of the fed-batch process. Data displayed by combining D0 and D6 (**A**) or displayed independently by day in culture (**B**). K-mean clustering was applied to data, enriched GO terms with biological process (BP) from each cluster are highlighted.

**Supplementary Figure 6: Gene Copy Number, mRNA Expression and Total Protein Abundances from Clonal Cells. A:** Gene copy number of heavy chain (HC) and light chain (LC) for each clone by ddPCR at D0. **B:** mRNA expression of heavy chain and light chain for each clone derived from RNA-seq data at D0. **C:** Total protein abundance of heavy chain and light chain for each clone derived from TMT-proteomics at D0. **D**: Correlation of protein abundance of light chain at D0 with the cell specific productivity (qP) from each clonal cell line.

**Supplementary Figure 7: Volcano Plots of Differentially Expressed Proteins Derived from TMT Proteomics. A-H:** Volcano plots of significant differentially expressed proteins from high secreting clones compared to low secreting clones expressing mAb1, MonoAb2, MonoAb1 and BisAb1 on D0 (**A-D**) and D6 (**E-H**) with significantly upregulated proteins in red and significantly downregulated proteins in blue. The size of the dot represents the number of peptides detected. Significance is defined as Log2FC > 2 and adjusted-P value (FDR) <0.05.

