## Supplementary Figure 1-7 for "Identification of Cellular Signatures Associated with Chinese Hamster Ovary (CHO) Cell Adaptation for Secretion of Antibodies"

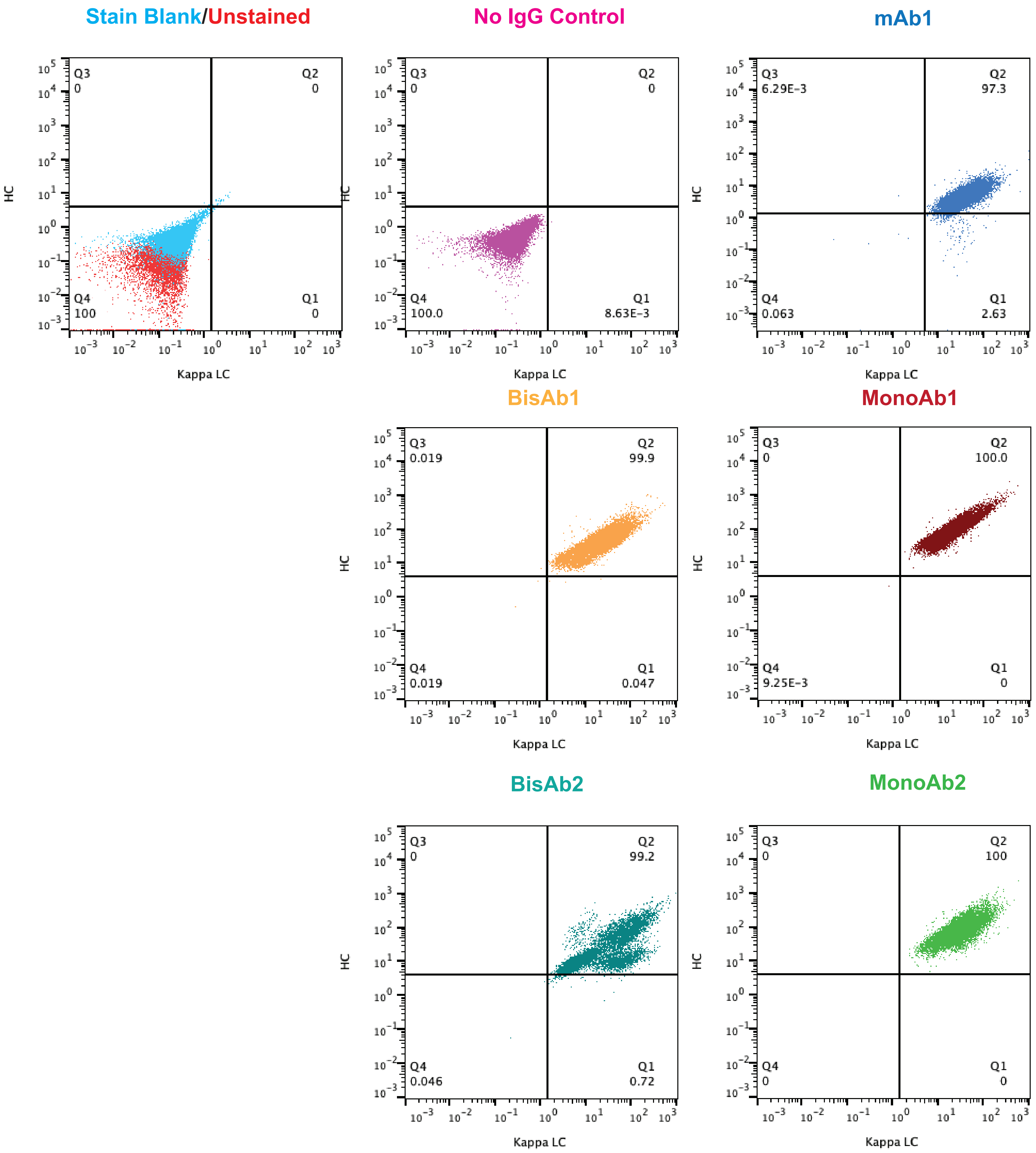

### Supplementary Figure 2

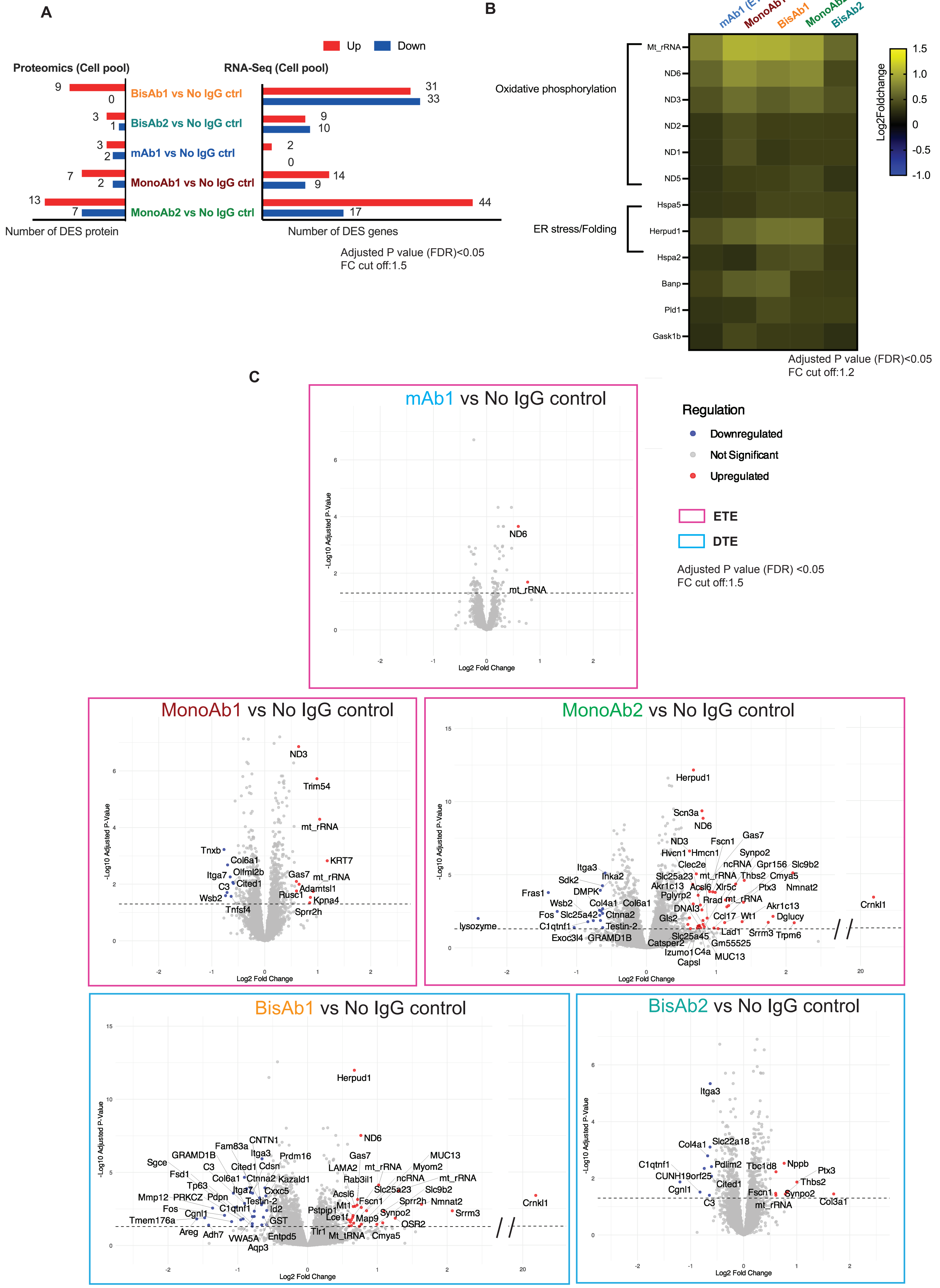

### Supplementary Figure 3

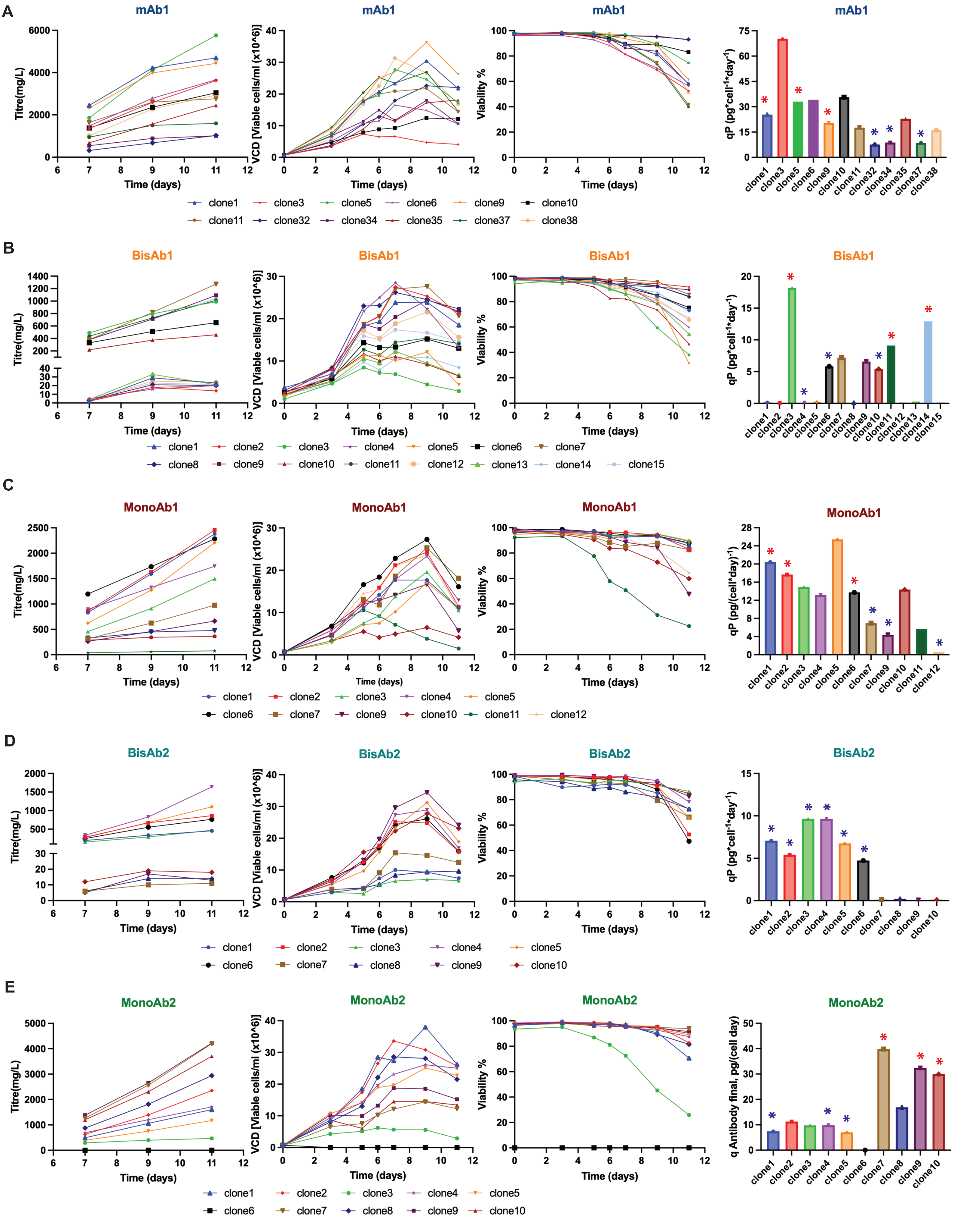

### Supplementary Figure 4

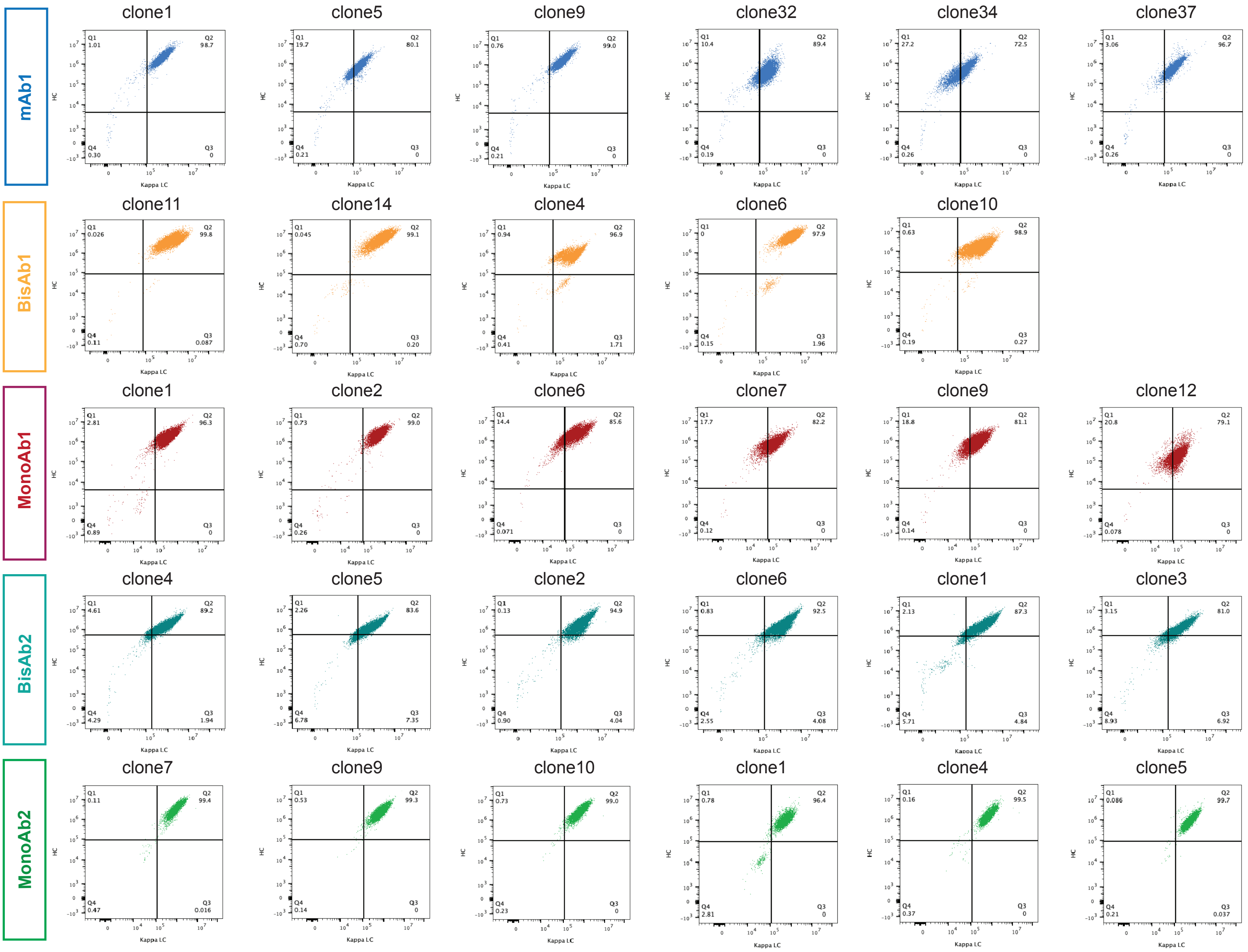

### Supplementary Figure 5

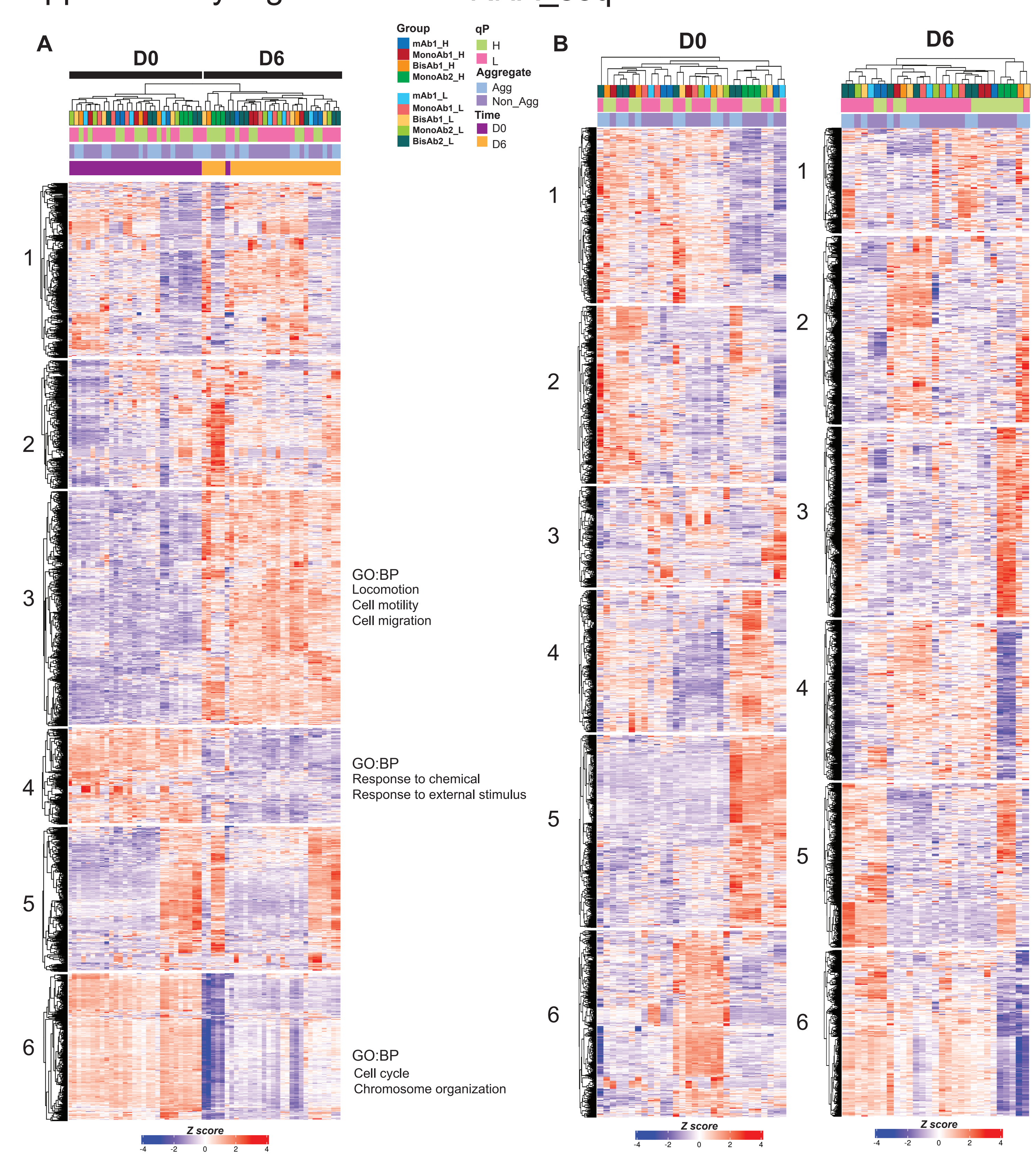

### Supplementary Figure 6

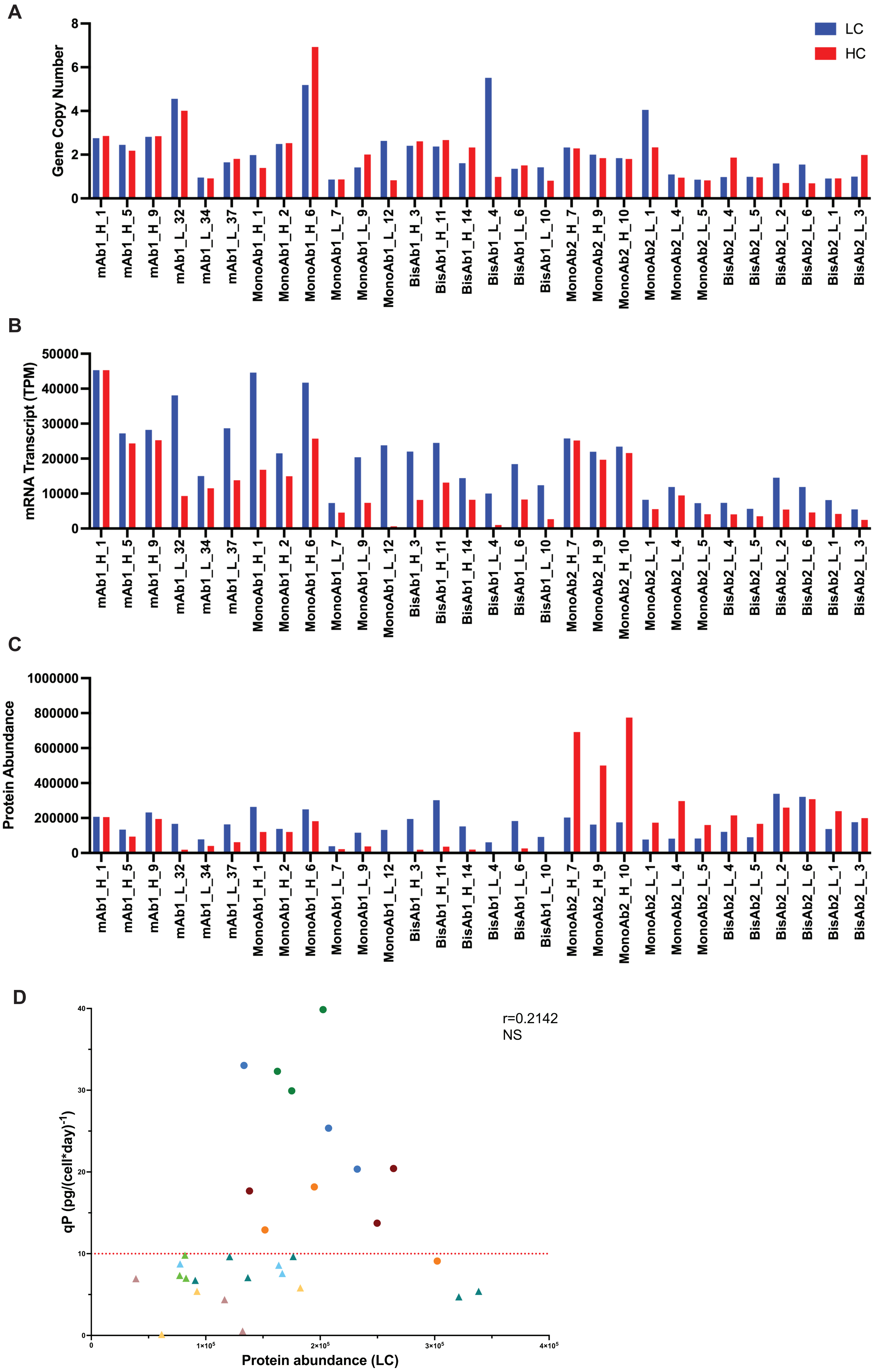

### Supplementary Figure 7

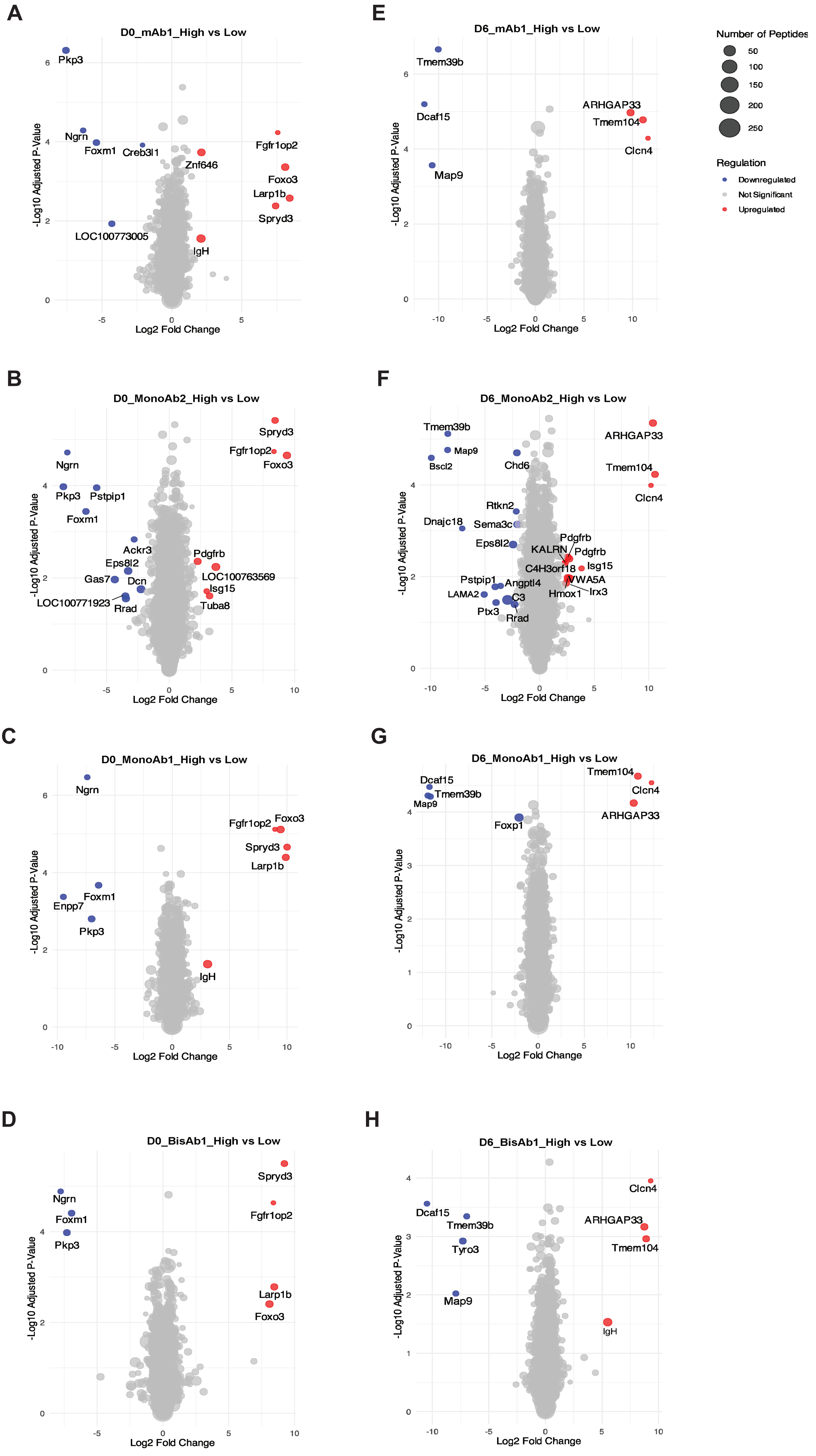
